# Scalable phylogenetic Gaussian process models improve the detectability of environmental signals on extinction risks for many Red List species

**DOI:** 10.1101/2023.06.21.545976

**Authors:** Misako Matsuba, Keita Fukasawa, Satoshi Aoki, Munemitsu Akasaka, Fumiko Ishihama

## Abstract

1. Conservation biologists have a daunting task of understanding the causes of species decline associated with anthropogenic factors and predicting the extinction risk of a growing number of endangered species. By stabilising estimates with information on closely related species, phylogenetic information among species can bridge gaps in information on species with small sample sizes when modelling large numbers of endangered species. However, modelling many species with the Gaussian process (GP), which underlies the evolutionary process of phylogenetic random effects, remains a challenge owing to the computational burden in estimating the large variance–covariance matrix.

2. Here, we applied a phylogenetic generalised mixed model with random slopes and random intercepts to 1,010 endangered vascular plant taxa in Japan following phylogenetic GPs implemented by nearest neighbour GP (NNGP) approximation. NNGP enables flexibility in changing the proximity on the phylogenetic tree of species from which information is borrowed to stabilise parameter estimates with a realistic computational burden. We evaluated the effectiveness of phylogenetic models by comparing the predictive performance and descriptive power of phylogenetic and non-phylogenetic models and identified the anthropogenic factors contributing to the decline of each of the studied endangered species.

3. We found that the model with phylogenetic information had better prediction performance than the model without phylogenetic information. The results showed that across all explanatory variables, the phylogenetic model could detect interspecific differences in response to environmental factors in a number of species more clearly. Combined with the phylogenetic signal results, we could also detect a phylogenetic bias in the species that could benefit from the positive effects of protected areas but reduce the extinction risk of 95% of all studied taxa.

4. In conclusion, our model, considering phylogenetic information with NNGP, allows the elucidation of factors causing the decline of many endangered species. In future analyses, the estimation of extinction probability linked to environmental change using such modelling might be applied to future climate–land use scenarios, advancing the comprehensive assessment of biodiversity degradation and threats to species at multiple scales.

## Introduction

Climate change and land cover change are major drivers of species extinction (Di Marco et al., 2019; Powers & Jetz, 2019). Current species extinction risks are already about 100–1,000 times higher than that in nature (Pimm et al., 2014), and the risk of biodiversity decline continues to increase (Butchart, 2010). Conservation biologists are now faced with the challenging task of reducing the extinction risk of the growing number of endangered species by elucidating the causes of species decline linked to environmental factors and predicting their future. However, Red List species often include species with extremely small population sizes and areas of occurrence (IUCN, 2001), which limits the identification of the factors underlying their decline and estimation of extinction risks linked to environmental change (Bachman et al., 2019; IUCN, 2001).

The availability of species’ phylogenetic information has been increasing in recent years (Beck et al., 2012; Mouquet et al., 2012), and it has the potential to improve extinction risk estimation of rare species. This is because branching patterns on evolutionary phylogenetic trees may help explain and predict interspecific correlates in biological and ecological processes, which are thought to reflect phenotypic, genetic, and behavioural differences among evolutionary lineages (Beck et al., 2012; Hernández et al., 2013). Especially in the field of macroecology, phylogenetic random effect models that incorporate species-specific responses to intrinsic and extrinsic factors correlated on a phylogenetic tree are considered powerful tools for multispecies systems because they can describe the likelihood of phylogenetically related species responding to an environmental driver in similar ways (Ives & Helmus, 2011; Li et al., 2020). However, modelling of evolutionary processes by Gaussian processes (GP) such as Wiener processes (Brownian motion) and Ornstein–Uhlenbeck (OU) processes, which are the underlying evolutionary processes of phylogenetic random effects, has not been put to practical use because of the huge computational load required to estimate a large variance–covariance matrix when assuming multiple species (Ives, 2018).

Nearest neighbour Gaussian process (NNGP) approximation is a scalable approach to GP model approximation with sparse representation (Datta et al., 2016a; Tikhonov et al., 2020) that has been developed in recent years in the field of spatial modelling. NNGP enables flexibility in the range of genetic distance correlation and hence in the range of closely related species from which we can borrow information to stabilise parameter estimates with a realistic computational burden. This is because NNGP uses a sparse precision matrix based on the nearest neighbour relationships among points to avoid the inverse computation of a huge variance–covariance matrix, which is a computational bottleneck in GP models (Datta et al., 2016a).

The objective of this study was to demonstrate the utility of applying a phylogenetic random effects model based on NNGP approximation in improving the estimation of extinction probabilities for endangered species, including many species with small sample sizes. The data used in the evaluation were the results of a comprehensive survey of 1,010 endangered vascular plant taxa across Japan, documenting changes in distribution over three time periods. By applying the phylogenetic random effects model to such spatiotemporally enriched data, we illustrated the first example of the strength of a model that utilises phylogenetic information to model a wide variety of endangered species.

## Material and Methods

### Data of threatened vascular plants in Japan

Data on threatened vascular plants were obtained from surveys conducted by the Japanese Society for Plant Systematics and the Ministry of the Environment for the preparation of the Red Data Book of Vascular Plants with the cooperation of volunteer surveyors from all over Japan. Surveys were conducted in three periods: 1994–1995 (hereafter written as’95), 2003–2004 (’04), and 2010–2011 (’11). The survey covered the entire country of Japan and was compiled at a spatial resolution of 5′ latitude and 7′ 30″ longitude (approximately 10 km grid). These data contain records of population sizes or events of extinction for each species classified as Near Threatened or higher. The population size was recorded by expert opinion, not by actual measurement. Because the focus of this study was to evaluate the effects of environmental factors on population viability over two time periods, we first extracted presence or extinction information for each population of a species as a response variable. In addition, population information from one period prior was extracted to account for the impact of population information. Thus, paired records for two periods, ‘95–’04 and ‘04–’11, could be compiled, with 1,010 taxa recorded from 2,113 taxa listed in the 2^nd^ to 4^th^ Red Data Book of Vascular Plants (1,010 in ‘95–’04 and 186 in ‘04–’11) and 9,623 pairs recorded (8,765 pairs in ‘95–’04, and 858 pairs in ‘04–’11). The 1,010 taxa contained 953 species, which further included 47 subspecies, 170 varieties, and 2 forma in 133 families. The average number of pairs recorded per species was 8.55, with 160.0 at maximum and 4.0 at median in ‘95–’04 and 4.60 and 52 at maximum and 2.0 at median in ‘04–’11, indicating that the data cover a wide variety of taxa with small samples.

### Phylogenetic data

Phylogenetic distance values for a pair of species were used for phylogenetic information on endangered vascular plants. To obtain the phylogenetic distance values of endangered vascular plants, their phylogenetic trees were generated using the phylo.maker function of the R package V.PhyloMaker (set as tree = GBOTB.extended, nodes = nodes.info.1, scenarios = “S1” Jin and Qian 2019). V.PhyloMaker used the updated and extended version of the dated megaphylogeny GBOTB reported by Smith & Brown (2018) as the backbone to generate phylogeny. Based on the generated phylogenetic trees, phylogenetic distances between species were calculated using the cophenetic.phylo function of the R package ape (Paradis et al., 2004). Specifically, the lengths of the branches of the phylogenetic tree were used to calculate the distance between pairs of phylogenetic tree tips. For the following analysis, the genetic distance was scaled so that the maximum distance equals to 2.0.

### Environmental data

We considered two climatic factors (mean annual temperature and annual precipitation) and seven land use factors (agricultural, urban, volcanic, wasteland, coastal, river/lake, and protected areas) as environmental factors affecting the extinction risk of threatened vascular plants based on a previous study (Watanabe et al., 2014). Average annual temperature and annual precipitation were calculated from daily data (10 km grid) of the Agro-Meteorological Grid Square Data, NARO (https://amu.rd.naro.go.jp/), for 2003 and 2010. For the percentages of agricultural land, urban areas, wastelands, coasts, river/lake, and forest area, and land use data (approximately 1 km grid) from the National Land Numerical Information were used to create each land use percentage on a 10 km grid unit (https://nlftp.mlit.go.jp /ksj/gml/datalist/KsjTmplt-L03-a.html). However, since the year of data release (1991, 1997, 2006, 2009, and 2014) did not match the year of the vascular plant survey, the published land use data were interpolated to one-year increments, and we obtained the data in 1994 and 2003. Inverse time-weighted interpolation was applied to the time series data (see Fujita et al. (2019) and Ohashi et al. (2019) for details on the calculation process). Among land uses, since the total area of agricultural land, artificial land, wasteland, coast, and river/lake together accounted for 100% of the total area, forest area, which has a large proportion, was excluded to allow the extraction of the effects of other land uses. All land uses included in the model were used as percentage values. For the volcanic area ratio, the raster data from the 1/200,000 land classification map compiled by the Ministry of Land, Infrastructure, Transport and Tourism was used to obtain the ratio in units of a 10 km grid. The survey was conducted from 1967 to 1978, and the topographic classifications such as volcanic lands have not changed enough to affect the 10 km grid calculation to this day. As for protected areas, the years of establishment of national parks and quasi-national parks in the National Land Survey Data differ for each park; thus, we extracted parks established before 1994 and 2003, respectively, and compiled the data for the two periods.

### Statistical models

In this study, we employed a phylogenetic generalised mixed model (PGLMM; Ives & Helmus, 2011) with random slopes and random intercepts following a scalable phylogenetic GP implemented by NNGP approximation (Datta et al., 2016a), which is a sparse and fast approximation for GP models. This approach can address situations wherein phylogenetically related species share common responses to an environmental factor. By considering phylogenetic correlation on the species-specific slopes and intercepts, it may be possible to estimate more robust species extinction probabilities with smaller estimation errors for many species, including species with small sample sizes, by utilising information of closely related species.

To estimate population-level extinction risks of Red List species and the effects of environmental factors on them, the presence and extinction of each population conditional on the population size of one period prior, which is known to affect the population viability (Chaudhary & Oli, 2020), was modelled using binomial PGLMM. Let *y_ijt_* denote survival (1) or extinction (0) of the population of species *i* at site *j* in year *t*; the binomial PGLMM with logit link function is as follows:

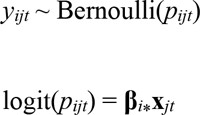

where **β***_i⁎_* is a vector of regression coefficients for species *i*, and **x***_jt_* is a design vector at site *j* in year *t*. The design vector consists of intercept, environmental factors (annual mean temperature, annual precipitation, agricultural land, urban area, volcanic land, wasteland, coast, river/lake, and protected areas), and population size class in the year of the previous survey (Matsuhashi et al., 2021). As the population size class is an ordered factor, we transformed it into a series of polynomial contrasts corresponding to linear, quadratic, and cubic trends (Chambers & Hastie, 2017).

The variation of regression coefficients among species are often partitioned into phylogenetically correlated and non-correlated variation in PGLMM (Ives & Helmus, 2011). In this study, we applied this two-part approach because it can accommodate a continuum between simple random variation and a fully phylogenetic structure. Considering the regression coefficient *m* for all the species, **β***_⁎m_* GP is as follows:

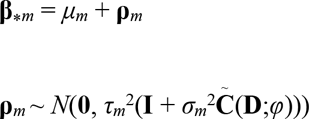

where *μ_m_* is mean of the *m*^th^ regression coefficient, **ρ***_m_* is the species-specific deviation from *μ_m_* subject to GP, *τ_m_*^2^ is the overall variation of random effects, and *σ_m_*^2^ is the conditional variance of phylogenetic components relative to structureless components. 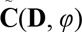 is the NNGP approximation of the covariance matrix **C**(**D**, *φ*), representing phylogenetic correlation depending on the matrix of genetic distances **D**. In this study, we applied the exponential correlation function **C**(**D**, *φ*) = exp(-*φ***D**), which corresponds to the OU process of trait evolution. To retain simplicity and model identifiability, we let *φ* be shared among covariates. We defined an indicator of phylogenetic signal as (conditional variance of phylogenetic component)/{(conditional variance of phylogenetic component) + (conditional variance of non-structured component)} = *σ_m_*^2^/(1 + *σ_m_*^2^) which ranges (0, 1).

Here, we summarise the formulation of NNGP. NNGP is a scalable approach for large geostatistical datasets that approximates GP with a sparse precision matrix 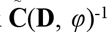 (Datta et al., 2016a, 2016b; Zhang et al., 2019). NNGP is based on the conditional independence of random vector **ρ** = (*ρ*_1_,…*ρ_n_*):

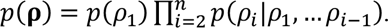

This formulation considers a directed acyclic graph (DAG) among *ρ_i_*s, whose directions of nodes are determined by an arbitrary ordering rule. In this study, we applied classic multidimensional scaling (MDS; Gower 1966) to the phylogenetic distance matrix and ordered species by the values of the first principal axis. Note that NNGP approximation is robust to the choice of ordering rule (Datta et al. 2016a). Assuming that **ρ** follows multivariate Gaussian distribution, *p*(*ρ_i_*|*ρ*_1_,…, *ρ_i_*_-1_) can be written as a linear model. Then, we can write a multivariate Gaussian density of **ρ** as a vector linear model:

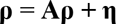

where **A** is *n* × *n* strictly lower-triangular matrix, and **η** is a vector of independent Gaussian random variables with mean **0** and diagonal covariance matrix **Λ**. **A** and **Λ** have an obvious relationship with the Cholesky decomposition of the covariance matrix **C**(**D**, *φ*) = **LΛL′** where **L** = (**I** – **A**)^-1^.

A major obstacle in estimating a GP model on a large dataset is the calculation of the Cholesky decomposition of the precision matrix for evaluating multivariate Gaussian density, which is *O*(*n*^3^) computational order. NNGP approximates the Cholesky decomposition of a precision matrix by replacing conditional probability *p*(*ρ_i_*|*ρ*_1_,…, *ρ_i_*_-1_) with *p*(*ρ_i_*|**ρ***_ν_*_(*I,k*)_), which is conditional on the *k* (<< *n*) nearest neighbours of the *i*^th^ sample on the DAG, *ν*(*i,k*). This approximation results in sparse **A** in which only the nearest neighbours of *i* have a non-zero value in the *i*^th^ row. Non-zero values of **A** in the *i*^th^ row, **A**[*i*, *ν*(*i,k*)], are determined by kriging weights based on nearest neighbours (Zhang et al., 2019):

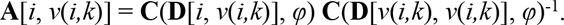

The diagonal element of **Λ**, **Λ**[*i, i*], is the kriging variance, which is variance conditional on the nearest neighbours, var(**ρ***_i_*| **ρ***_ν_*_(*i,k*)_), as follows:

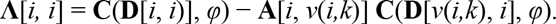

Then, precision matrix 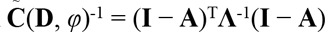 is also sparse. In this study, we set *k* = 5, which is as large as possible in terms of computational time.

We estimated parameters of the NNGP-PGLMM by Bayesian inference via MCMC sampling. Prior setting is essential for successful prediction by the GP model, because hyperparameters of GP are often unidentifiable and have non-regular geometry of the joint posterior (Zhang, 2004). Especially, range parameters that are too small (i.e. inverse of *φ*) often degenerate the covariance function to Kronecker’s delta, resulting in the redundancy of the non-structured and structured components of random effects. To avoid model redundancy, we applied a boundary-avoiding prior (Gelman et al., 2014) on the range parameter. We reparametrised *φ* to a range parameter as 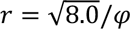 corresponding to the scaled genetic distance that gives a correlation of 0.06 and set inverse gamma prior IG (10, 10) on *r* whose 95% range is (0.585, 2.08). This density function is positive-valued and convex around 0, which can penalise values of correlation range that are too small. Although posterior of hyperparameters such as range of covariance function can be affected by the prior settings, it is known that prediction by GP is quite robust (Chen & Wang, 2016; Zhang, 2004). We applied weakly informative or vague prior for the other parameters: *μ_m_ ∼ N*(0, 1)*, τ_m_ ∼ half-N*(0, 1) and *σ_m_* ∼ *half-Cauchy*(0, 1). The prior of *σ_m_* implies no prior information about the phylogenetic signal because *σ_m_*^2^/(1 + *σ_m_*^2^) ∼ Beta(0.5, 0.5), which is a symmetric U-shaped distribution.

We obtained posterior samples by No-U-Turn sampler using stan 2.29.2 (Stan Development Team 2022). We sampled by four MCMC chains with 10,000 iterations after discarding the first 3,000 as burn-in. To reduce memory usage, we took one sample per 10 iterations and 4,000 posterior samples for inference. We checked the mixing of multiple chains using Rhat (Gelman & Rubin 1992) and visual inspection of trace plots. To see how much the prediction performance of the model was improved by considering phylogenetic relationships, we compared the Watanabe–Akaike information criterion (WAIC; Watanabe, 2010) and leave-one-out cross-validation (LOO; Vehtari et al. 2017) with the null model of phylogenetic signals in which **ρ***_m_* ∼ *N*(**0**, *τ_m_*^2^**I**). To evaluate the descriptive power of the phylogenetic model, the area under curve (AUC; Swets, 1988) for training data was compared to the null model of the phylogenetic model.

The model without considering phylogenetic relationships was derived by a coefficient **β** that did not include the phylogenetic random effect **ρ**_*m*_, but only the random effect **ε**_*m*_ without structure, as shown below. We sampled by four MCMC chains with 10,000 iterations after discarding the first 1,000 as burn-in.

## Results

All chains converged adequately in all models (mean Rhat < 1.05). The results indicated that the model with phylogenetic information had better prediction performance than the model without phylogenetic information, since both WAIC and LOO values were smaller in the model with phylogenetic information than in the model without phylogenetic information (phylogenetic model:non-phylogenetic model = 4227.8:4230.5 in WAIC, 4245.5:4247.7 in LOO). The phylogenetic signals confirmed that phylogenetic effects were significant, especially for the coefficient of protected areas (Fig. 3; phylogenetic signal = 0.54 ± 0.28 for protected areas and 0.34 ± 0.30 for the average of all variables). The AUC values were comparable between the two models (phylogenetic model:non-phylogenetic model = 0.911:0.913), indicating that the descriptive power of the present phylogenetic model is comparable.

Among the 10 fixed factors in the model, the 95% confidence interval (CI) of coefficients of four environmental factors (protected areas, proportion of wasteland, artificial land, and agricultural land) and the population size class from one period prior did not overlap zero (Fig. 1; Appendix B). The 95% CI of coefficients of the other factors tested (mean annual temperature, annual precipitation, volcanic land, proportion of coastal area, and river/lake) overlapped with zero. The four influencing environmental variables were revealed to have a positive contribution from the proportion of wasteland and protected areas, and a negative influence from the proportion of urban area and agricultural land (Fig. 1). Furthermore, the number of species affected was 960 (95% of the total number of taxa analysed) in protected areas, 960 in the proportion of wasteland, 963 in the proportion of agricultural land, and 866 in the proportion of artificial land.

**Fig. 1.**
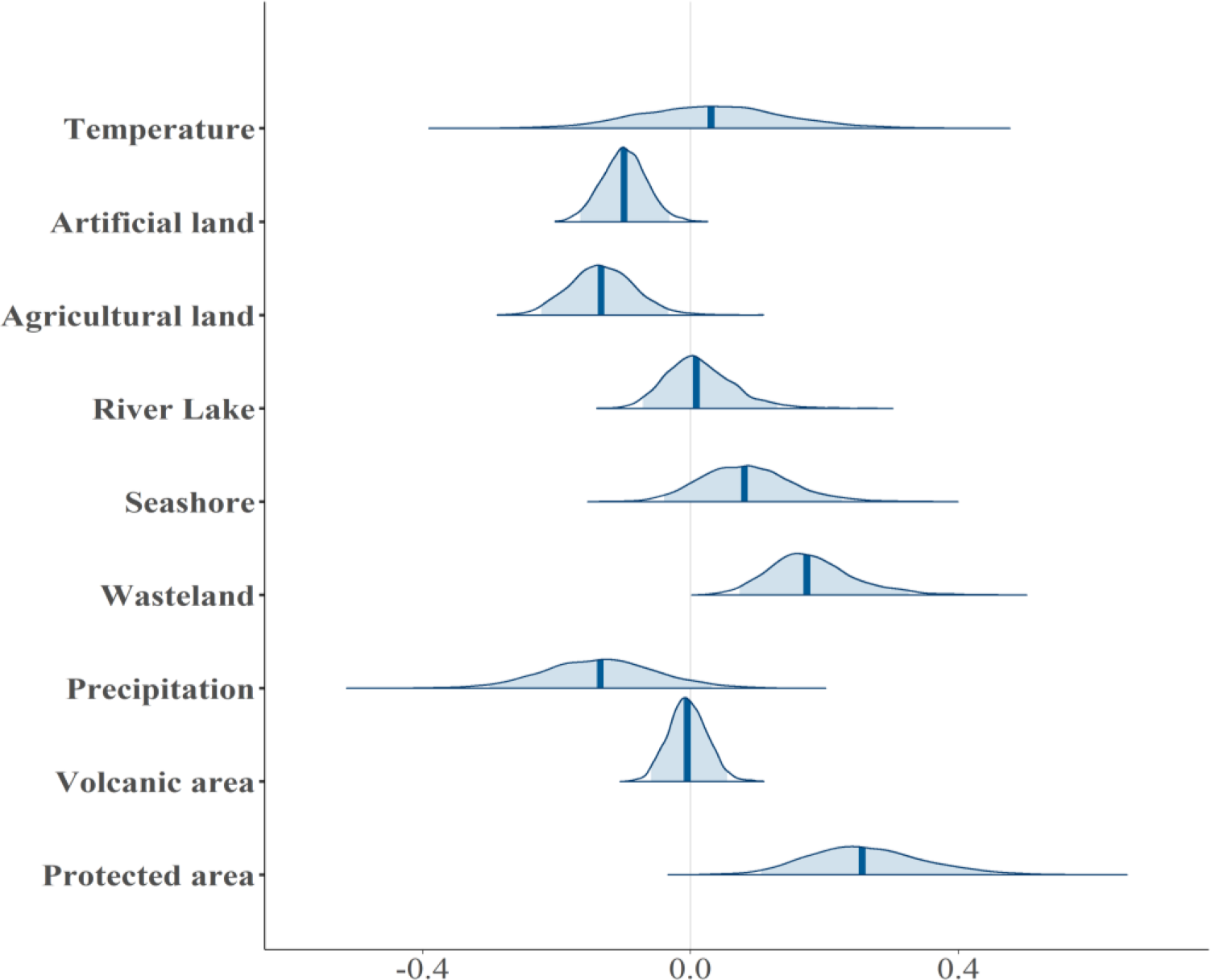
Average values of posterior distributions with medians and 95% CI intervals for each environmental variable in the model.

The results of the comparison of χ^2^ values for each species showed that for many species, we could detect interspecific differences in response to environmental factors more clearly in the phylogenetic model throughout all explanatory variables (Fig. 2). Even species that showed values near zero in the model without phylogenetic information were shown to have much larger values (e.g. Protected areas; Fig. 2) when phylogenetic information was considered. Among species for which the clear effect of environments was detected because 95% CI did not overlap with zero, fewer species had reversed positive or negative signs of the estimates between phylogenetic and non-phylogenetic models (the maximum and minimum percentage of species that had reversed numbers were 20.6% for temperature and 0% for the proportion of artificial land, agricultural land, wasteland, and seashore).

**Fig. 2.**
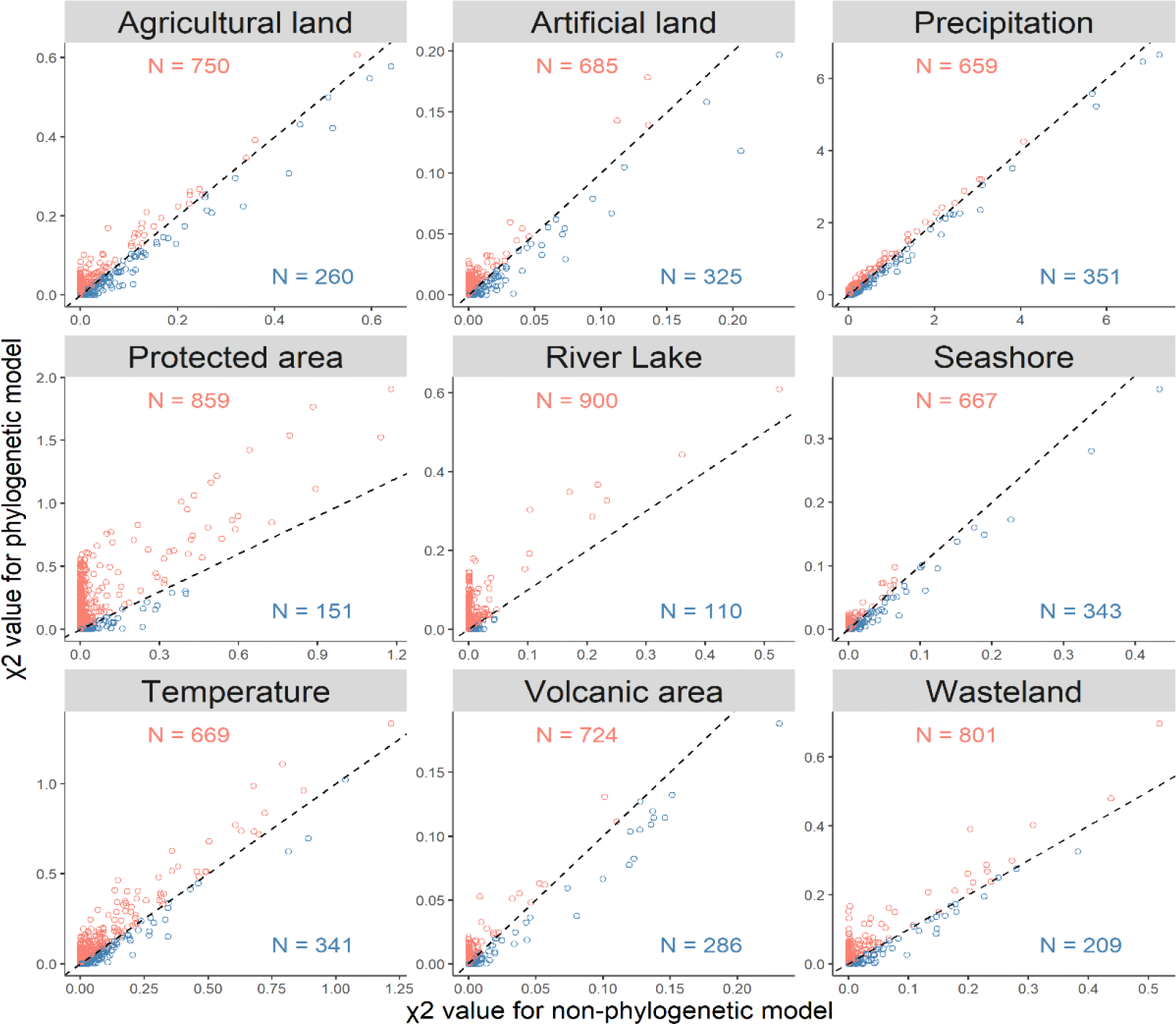
Results of the comparison of χ^2^ values of each species for each environmental variable.

**Fig. 3.**
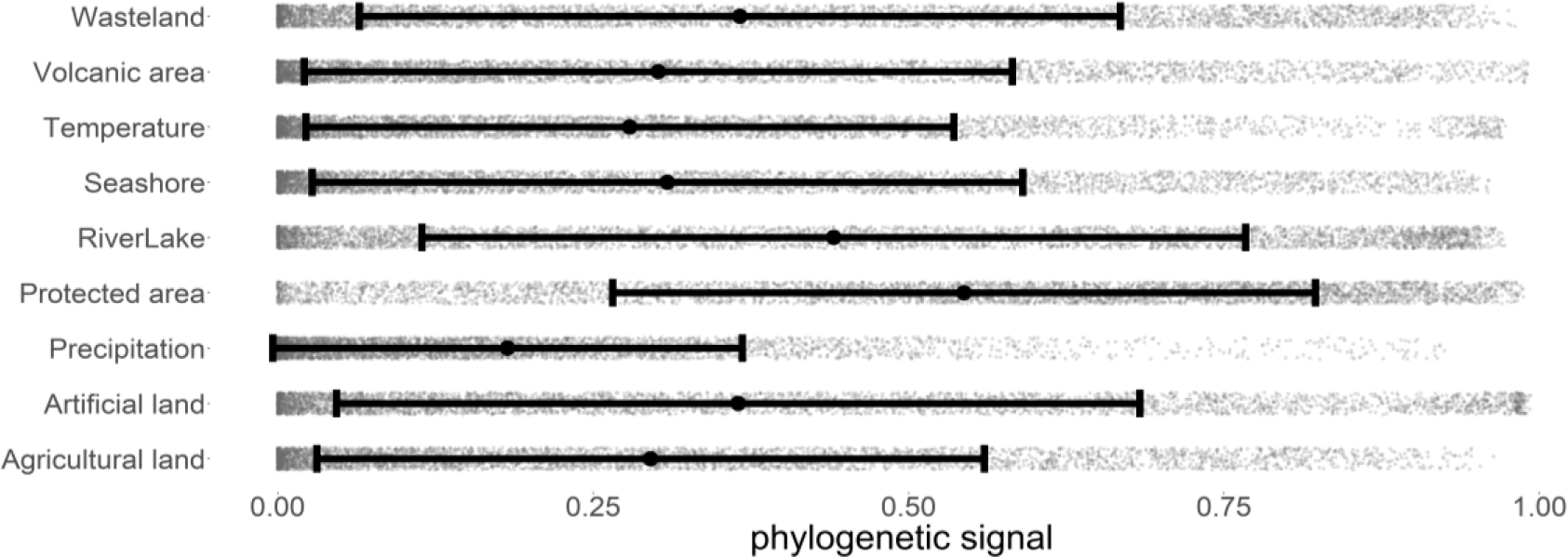
Results of phylogenetic signals of each environmental variable.

Figure 4 shows the extent to which the coefficients of the explanatory variables varied phylogenetically. The results showed that the explanatory variables reducing extinction risk of endangered vascular plants had a phylogenetic cluster structure, as well as positive and negative differences in their estimates. This indicates that the effects of environmental factors are phylogenetically dependent. The phylogenetic half-life (the amount of time expected for a trait to move halfway to the mean value; Hansen 1997) was 0.266 for posterior distributions and 0.271 for prior distributions (Appendix A).

**Fig. 4a.**
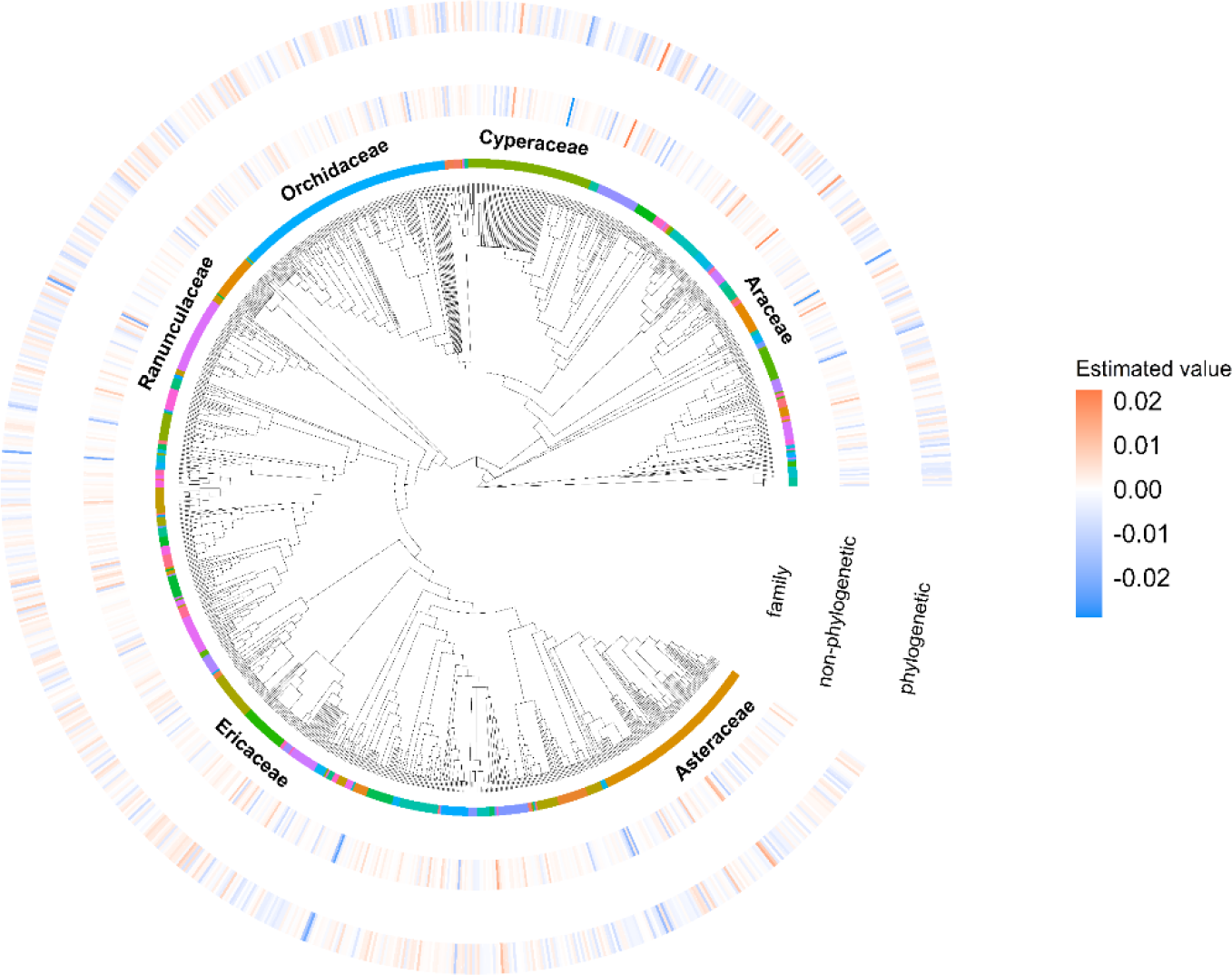
Comparison of estimated coefficients of artificial land between phylogenetic model and non-phylogenetic model.

**Fig. 4b.**
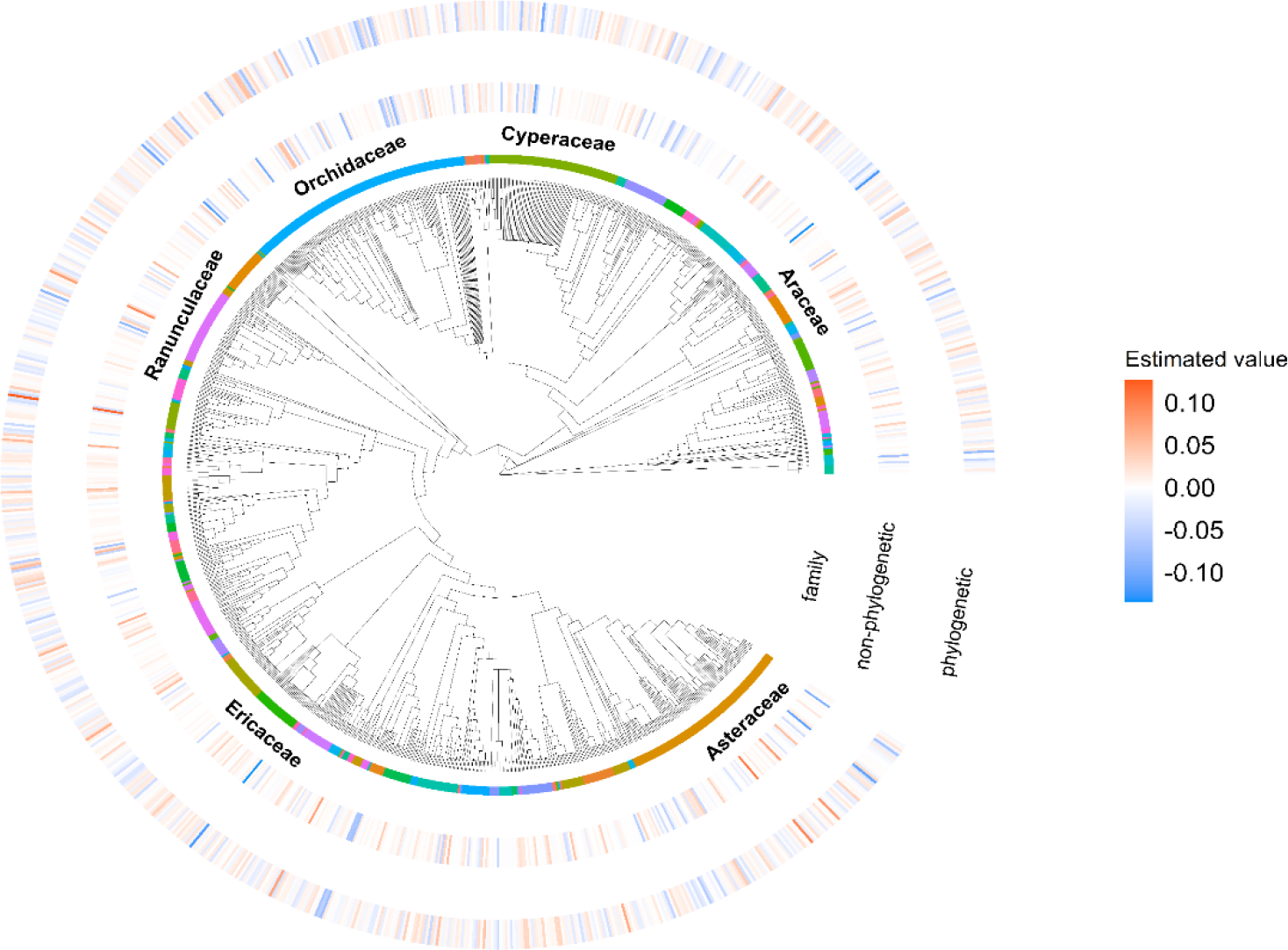
Comparison of estimated coefficients of agricultural land between phylogenetic model and non-phylogenetic model.

**Fig. 4c.**
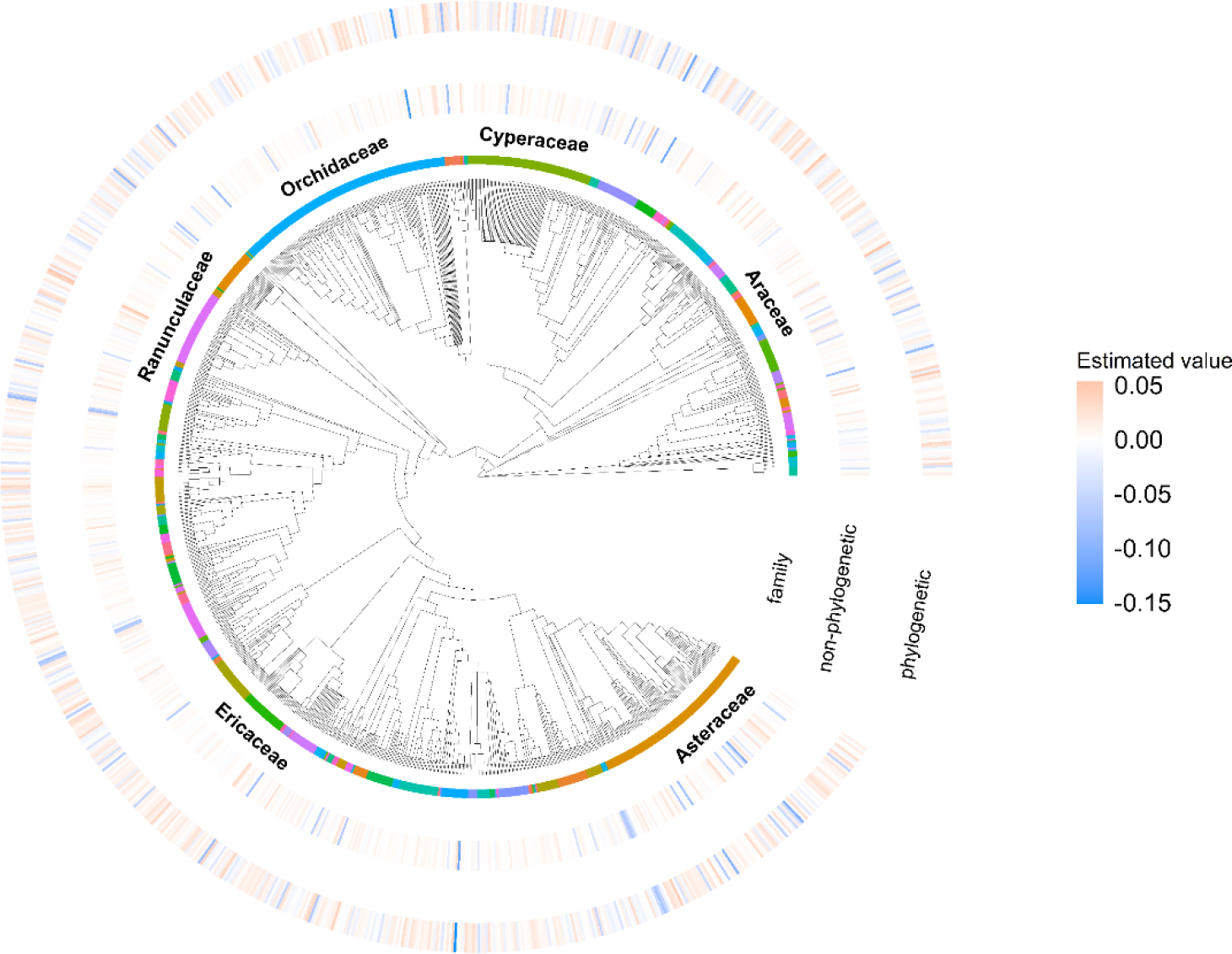
Comparison of estimated coefficients of wasteland between phylogenetic model and non-phylogenetic model.

**Fig. 4d.**
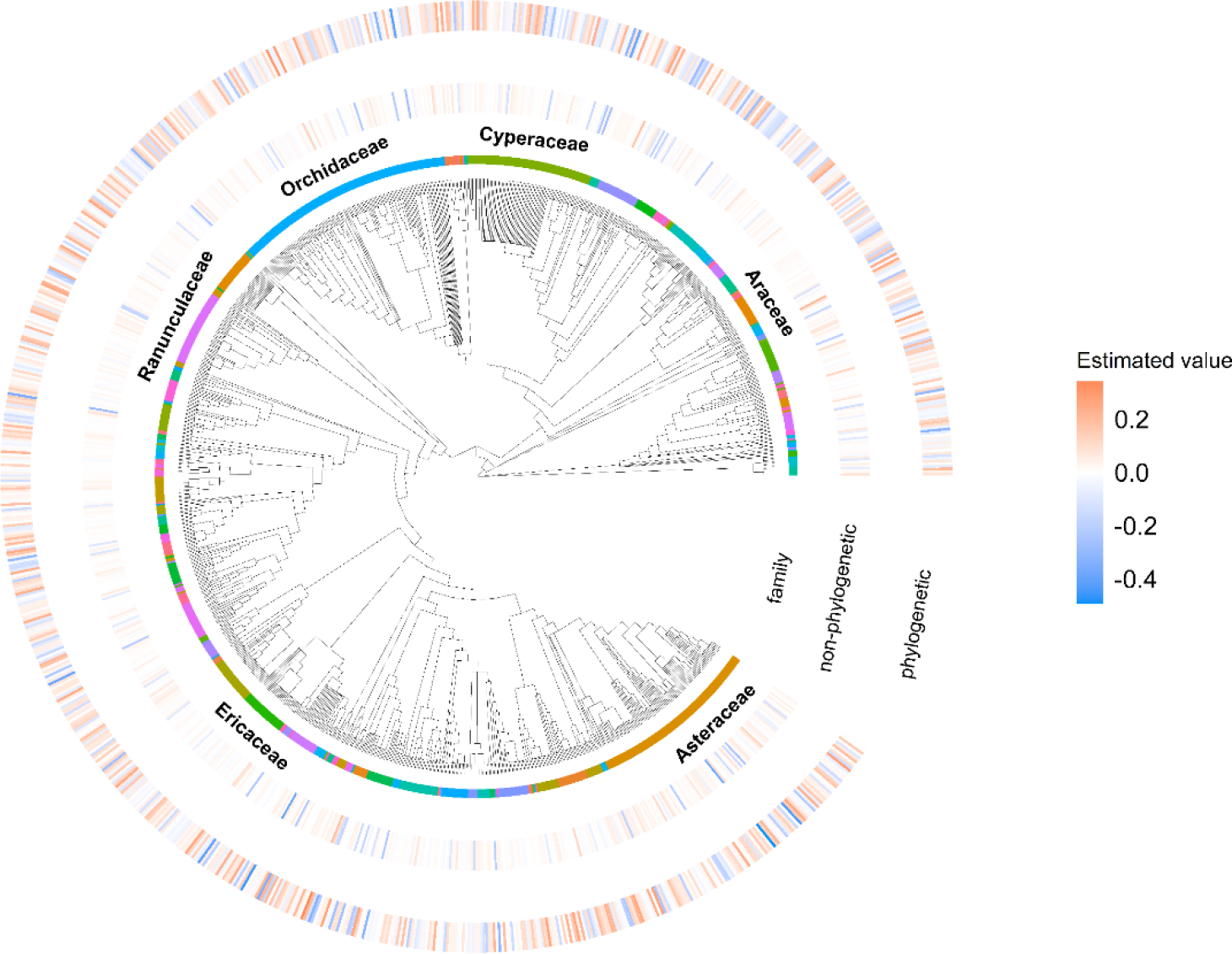
Comparison of estimated coefficients of protected areas between phylogenetic model and non-phylogenetic model.

## Discussion

We demonstrated that application of the NNGP approximation to 1,010 endangered vascular plant species, taking into account species phylogenetic information, improved the detection of species-specific responses to the environment without compromising prediction accuracy. In particular, the present data contained a large number of species (238 species, 24%) with a small sample size of fewer than 10 pairs of records. Previous studies on estimating the extinction risk of endangered species have found it difficult to address such small samples (Bachman et al., 2019; Walker et al., 2020), often excluding many species with limited data and analysing a single species or a few species with abundant data (Bachman et al., 2019; IUCN, 2001; Walker et al., 2020). In this study, we demonstrated that such a challenge can be resolved by using an approximation of the GP model, based on an evolutionary model of the OU process, to achieve a comprehensive analysis of multiple species with a wide range of sample sizes. Although this study focused on endangered species as a representative example of a problem with a small sample size, data obtained in ecosystems that include species and fields that are difficult to survey (e.g. deep sea ecosystem) could also benefit from the application of this model, and it is expected that practical use will expand beyond endangered species in the future.

When modelling multiple species concurrently, the incorporation of the evolutionary background is thought to have advantages, such as stabilising estimates by borrowing information from closely related species (Cooper et al., 2016; Münkemüller et al., 2015), although whether niche evolutionary processes can be inferred from species occurrence patterns has been controversial. Although the calculated phylogenetic signal is conditional on model structure and prior distribution, and thus may not be expected to provide accurate inferences about evolutionary processes, our results indicate that phylogenetic structure is a useful measure in assessing extinction risk. Again, our approach highlights the practical benefits of considering phylogeny in multispecies extinction risk assessments. It is known that the GP model used in this study has high prediction accuracy in spatial statistics (Golding & Purse, 2016), but the large computational load poses a problem (Ives, 2018). The present approach overcomes this limitation by utilising the recently developed NNGP model approximation (Datta et al., 2016a; Tikhonov et al., 2020) in ecology, allowing more flexible phylogenetic models to be applied to a wide variety of species. For the flexibility of the model, we used the covariance of the OU process, which is a common exponential model In spatial statistics, and this process seemed to have the same advantages in systematic modelling as spatial smoothing. The OU process is a generalisation of the Brownian motion model, and it is possible to determine the extent to which information is borrowed according to the posterior probability maximisation criteria. On the other hand, the problem of weak identifiability of hyperparameters and the associated poor geometry of the posterior distribution needs to be addressed (Zhang, 2004). In overcoming such obstacles, it is beneficial to introduce an appropriate prior distribution in the Bayesian modelling framework.

The model results indicated that the establishment of protected areas and land use modifications affect the presence and extinction of species. Combined with the results of the phylogenetic signal, we showed that there was a phylogenetic bias in the species that could benefit from the positive effects of protected areas (coefficient in phylogenetic model:non-phylogenetic model = 0.26 ± 0.08:0.25 ± 0.09).

This means that there is a phylogenetic correlation in the sensitivity to various decline pressures (development, exploitation, etc.) that are mitigated by the establishment of protected areas and conservation measures (Akasaka et al., 2017; Kadoya et al., 2014). On the other hand, we did not find strong phylogenetic correlations for artificial land, agricultural land, and wastelands. This is consistent with the point that urban expansion has long had a devastating impact on the extinction of various species (Czech et al., 2000; McKinney, 2006). It is likely that no trend was observed in agricultural lands because they are mosaic environments with a variety of species with different life history traits (Bennett et al., 2006; Graham et al., 2019; Sugimoto et al., 2022). Similarly, wastelands contain various types of secondary grassland and shoreline vegetation (Akasaka et al., 2014), which may have affected various species in a phylogenetic manner.

In conclusion, the model presented in this study, which leverages phylogenetic information, has made it possible to elucidate the factors causing the decline of a number of endangered species, which was previously difficult to achieve. Coupling the model with climatic and land use factors will enable the prediction of the future extinction risk or of the population size of endangered species. These predictions can then be analysed in complementary analyses to provide powerful information for selecting conservation priority areas (Pressey et al., 2007). Such an approach is likely to become even more important today, when conservation resources are limited both economically and in terms of human resources, warranting efficient conservation efforts (Butchart et al., 2015). We would also like to mention that explicit consideration of phylogenetic structure is a key advantage of this modelling approach as it allows for the discussion of phylogenetic diversity and the decline in ecosystem function. In future analyses, it is expected that the estimation of extinction probability linked to environmental change enabled by this modelling will be applied to future climate–land use scenarios. This will enhance the comprehensive assessment of biodiversity degradation at multiple scales—population, species, community—encompassing endangered species.

## AUTHOR CONTRIBUTIONS

M.M., K.F., and F.I. conceived the idea and designed the methodology of the study; M.M. and F.I. collected the data; M.M. and K.F. analysed the data; M.M., K.F. and F.I. led the writing of the manuscript; all authors contributed critically to the drafts and gave final approval for publication.

## Supporting information

coefficients

## ACKNOWLEDGEMENT

We thank Tagane S. for comments on this manuscript. We would like to express our gratitude to the Japanese Society for Plant Systematics for providing the vascular plant distribution data.

---

**Appendix A.**
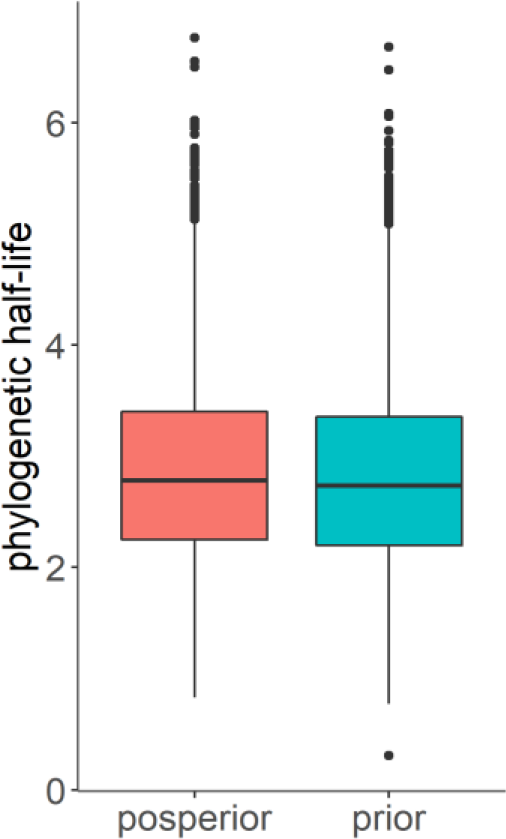
Results of phylogenetic half-life for posterior and prior distributions.

**Appendix B.**
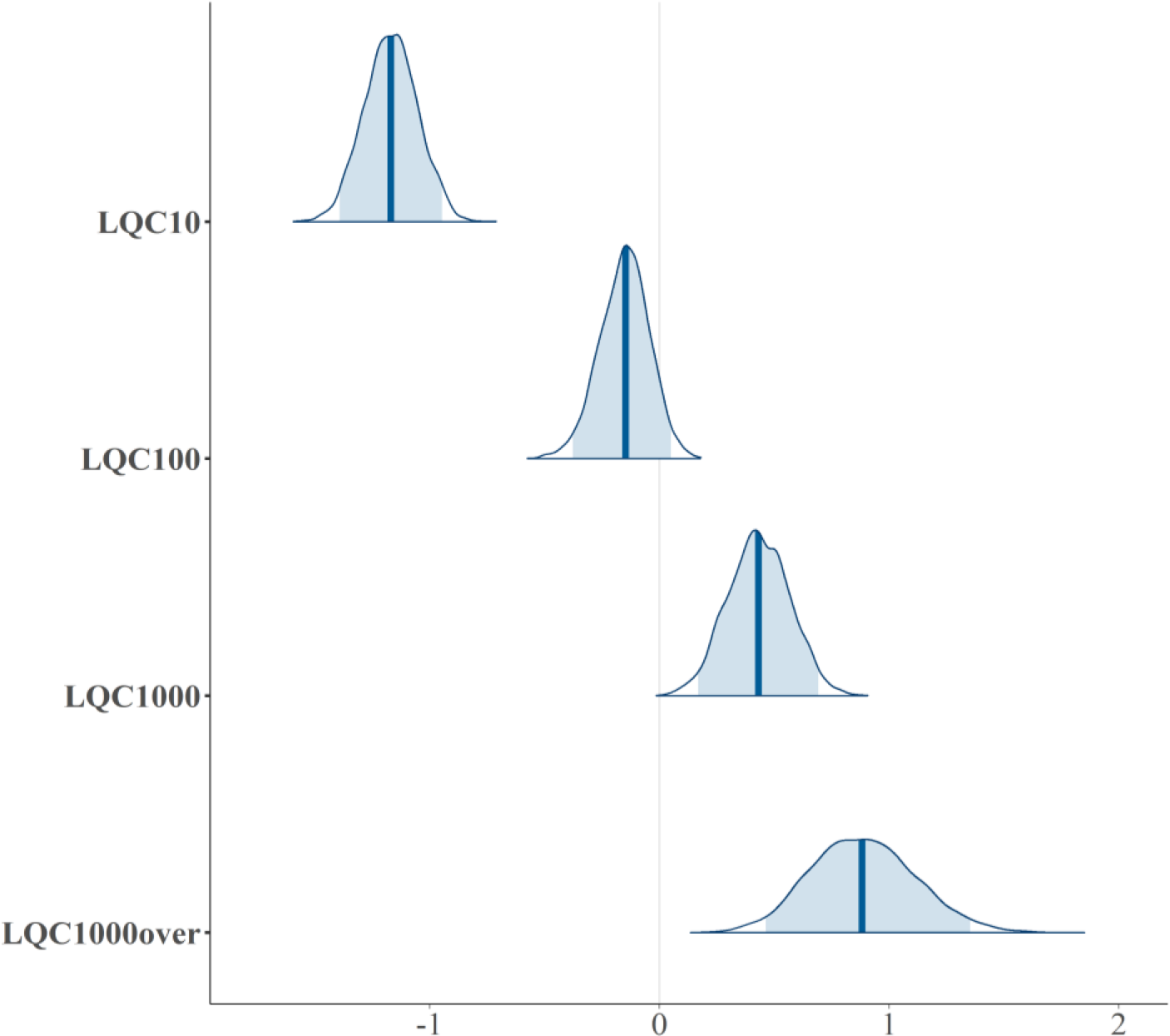
The average values of posterior distributions with medians and 95% CI intervals for each population size class from one period prior in the model.

## REFERENCES

Akasaka, M., Takenaka, A., Ishihama, F., Kadoya, T., Ogawa, M., Osawa, T., Yamakita, T., Tagane, S., Ishii, R., Nagai, S., Taki, H., Akasaka, T., Oguma, H., Suzuki, T., & Yamano, H. (2014). Development of a national land-use/cover dataset to estimate biodiversity and ecosystem services. In: S. Nakano, T. Yahara, T. Nakashizuka (Eds.) Integrative observations and assessments. Ecological research monographs (pp. 209–229). Springer, Tokyo.

Akasaka, M., Kadoya, T., Ishihama, F., Fujita, T., & Fuller, R. A. (2017). Smart protected area placement decelerates biodiversity loss: A representation-extinction feedback leads rare species to extinction. Conservation Letters, 10(5), 539–546. https://doi.org/10.1111/conl.12302

Bachman, S. P., Field, R., Reader, T., Raimondo, D., Donaldson, J., Schatz, G. E., & Lughadha, E. N. (2019). Progress, challenges and opportunities for Red Listing. Biological Conservation, 234, 45–55. https://doi.org/10.1016/j.biocon.2019.03.002

Beck, J., Ballesteros-Mejia, L., Buchmann, C. M., Dengler, J., Fritz, S. A., Gruber, B., Hof, C., Jansen, F., Knapp, S., Kreft, H., Schneider, A. K., Winter, M., & Dormann, C. F. (2012). What’s on the horizon for macroecology? Ecography, 35(8), 673–683. https://doi.org/10.1111/j.1600-0587.2012.07364.x

Bennett, A. F., Radford, J. Q., & Haslem, A. (2006). Properties of land mosaics: Implications for nature conservation in agricultural environments. Biological Conservation, 133(2), 250–264. https://doi.org/10.1016/j.biocon.2006.06.008

Butchart, S. H. M., Walpole, M., Collen, B., van Strien, A., Scharlemann, J. P. W., Almond, R. E. A., Baillie, J. E. M., Bomhard, B., Brown, C., Bruno, J., Carpenter, K. E., Carr, G. M., Chanson, J., Chenery, A. M., Csirke, J., Davidson, N. C., Dentener, F., Foster, M., Galli, A., Galloway, J. N.,…Watson, R. (2010). Global biodiversity: Indicators of recent declines. Science, 328(5982), 1164–1168. https://doi.org/10.1126/science.1187512

Butchart, S. H. M., Clarke, M., Smith, R. J., Sykes, R. E., Scharlemann, J. P. W., Harfoot, M., Buchanan, G. M., Angulo, A., Balmford, A., Bertzky, B., Brooks, T. M., Carpenter, K. E., Comeros-Raynal, M. T., Cornell, J., Ficetola, G. F., Fishpool, L. D. C., Fuller, R. A., Geldmann, J., Harwell, H.,…Burgess, N. D. (2015). Shortfalls and solutions for meeting national and global conservation area targets. Conservation Letters, 8(5), 329–337. https://doi.org/10.1111/conl.12158

Chambers, J. M., & Hastie, T. J. (2017). Statistical models. In Statistical models in S. Champman & Hall/CRC, London.

Chaudhary, V., & Oli, M. K. (2020). A critical appraisal of population viability analysis. Conservation Biology, 34(1), 26–40. https://doi.org/10.1111/cobi.13414

Chen, Z., & Wang, B. (2018). How priors of initial hyperparameters affect Gaussian process regression models. Neurocomputing, 275, 1702–1710. https://doi.org/10.1016/j.neucom.2017.10.028

Cooper, N., Thomas, G. H., Venditti, C., Meade, A., & Freckleton, R. P. (2016). A cautionary note on the use of Ornstein Uhlenbeck models in macroevolutionary studies. Biological Journal of the Linnean Society, 118(1), 64–77. https://doi.org/10.1111/bij.12701

Czech, B., Krausman, P. R., & Devers, P. K. (2000). Economic associations among causes of species endangerment in the United States. BioScience, 50(7), 593–601. https://doi.org/10.1641/0006-3568(2000)050[0593:EAACOS]2.0.CO;2

Datta, A., Banerjee, S., Finley, A. O., & Gelfand, A. E. (2016a). Hierarchical nearest-neighbor Gaussian process models for large geostatistical datasets. Journal of the American Statistical Association, 111(514), 800–812. https://doi.org/10.1080/01621459.2015.1044091

Datta, A., Banerjee, S., Finley, A. O., & Gelfand, A. E. (2016b). On nearest-neighbor Gaussian process models for massive spatial data. WIREs Computational Statistics, 8(5), 162–171. https://doi.org/10.1002/wics.1383

Di Marco, M., Venter, O., Possingham, H. P., & Watson, J. E. M. (2018). Changes in human footprint drive changes in species extinction risk. Nature Communications, 9(1), 4621. https://doi.org/10.1038/s41467-018-07049-5

Di Marco, M., Ferrier, S., Harwood, T. D., Hoskins, A. J., & Watson, J. E. M. (2019). Wilderness areas halve the extinction risk of terrestrial biodiversity. Nature, 573(7775), 582–585. https://doi.org/10.1038/s41586-019-1567-7

Fujita, T., Ariga, T., Ohashi, H., Hijioka, Y., & Fukasawa, K. (2019). Assessing the potential impacts of climate and population change on land-use changes projected to 2100 in Japan. Climate Research, 79(2), 139–149. https://doi.org/10.3354/cr01580

Gelman, A., Carlin, J. B., Stern, H. S., Dunson, D. B., Vehtari, A., & Rubin, D. B. (2014). Bayesian data analysis, third edition. Taylor & Francis.

Golding, N., & Purse, B. V. (2016). Fast and flexible Bayesian species distribution modelling using Gaussian processes. Methods in Ecology and Evolution, 7(5), 598–608. https://doi.org/10.1111/2041-210X.12523

Gower, J. C. (1966). Some distance properties of latent root and vector methods used in multivariate analysis. Biometrika, 53(3-4), 325–338. https://doi.org/10.1093/biomet/53.3-4.325

Graham, S. I., Kinnaird, M. F., O’Brien, T. G., Vågen, T. G., Winowiecki, L. A., Young, T. P., & Young, H. S. (2019). Effects of land-use change on community diversity and composition are highly variable among functional groups. Ecological Applications, 29(7), e01973. https://doi.org/10.1002/eap.1973

Hernández, C. E., Rodríguez-Serrano, E., Avaria-Llautureo, J., Inostroza-Michael, O., Morales-Pallero, B., Boric-Bargetto, D., Canales-Aguirre, C. B., Marquet, P. A., & Meade, A. (2013). Using phylogenetic information and the comparative method to evaluate hypotheses in macroecology. Methods in Ecology and Evolution, 4(5), 401–415. https://doi.org/10.1111/2041-210X.12033

IUCN. (2001). IUCN Red List categories and criteria: version 3.1. Prepared by the IUCN Species Survival Commission.

Ives, A. R. (2018). Mixed and phylogenetic models: A conceptual introduction to correlated data. Leanpub. https://leanpub.com/correlateddata

Ives, A. R., & Helmus, M. R. (2011). Generalized linear mixed models for phylogenetic analyses of community structure. Ecological Monographs, 81(3), 511–525. https://doi.org/10.1890/10-1264.1

Kadoya, T., Takenaka, A., Ishihama, F., Fujita, T., Ogawa, M., Katsuyama, T., Kadono, Y., Kawakubo, N., Serizawa, S., Takahashi, H., Takamiya, M., Fujii, S., Matsuda, H., Muneda, K., Yokota, M., Yonekura, K., & Yahara, T. (2014). Crisis of Japanese vascular flora shown by quantifying extinction risks for 1618 taxa. PLoS ONE, 9(6), e102384. https://doi.org/10.1371/journal.pone.0098954

Li, D., Dinnage, R., Nell, L. A., Helmus, M. R., & Ives, A. R. (2020). phyr: An R package for phylogenetic species-distribution modelling in ecological communities. Methods in Ecology and Evolution, 11(11), 1455–1463. https://doi.org/10.1111/2041-210X.13471

Matsuhashi, S., Asai, M., & Fukasawa, K. (2021). Estimations and projections of *Avena fatua* dynamics under multiple management scenarios in crop fields using simplified longitudinal monitoring. PLOS ONE, 16(1), e0245217. https://doi.org/10.1371/journal.pone.0245217

McKinney, M. L. (2006). Urbanization as a major cause of biotic homogenization. Biological Conservation, 127(3), 247–260. https://doi.org/10.1016/j.biocon.2005.09.005

Mouquet, N., Devictor, V., Meynard, C. N., Munoz, F., Bersier, L.-F., Chave, J., Couteron, P., Dalecky, A., Fontaine, C., Gravel, D., Hardy, O. J., Jabot, F., Lavergne, S., Leibold, M., Mouillot, D., Münkemüller, T., Pavoine, S., Prinzing, A., Rodrigues, A. S. L.,…Thuiller, W. (2012). Ecophylogenetics: Advances and perspectives. Biological Reviews, 87(4), 769–785. https://doi.org/10.1111/j.1469-185X.2012.00224.x

Münkemüller, T., Boucher, F. C., Thuiller, W., & Lavergne, S. (2015). Phylogenetic niche conservatism – Common pitfalls and ways forward. Functional Ecology, 29(5), 627–639. https://doi.org/10.1111/1365-2435.12388

Ohashi, H., Fukasawa, K., Ariga, T., Matsui, T., & Hijioka, Y. (2019). High-resolution national land use scenarios under a shrinking population in Japan. Transactions in GIS, 23(4), 786–804. https://doi.org/10.1111/tgis.12525

Paradis, E., Claude, J., & Strimmer, K. (2004). APE: Analyses of phylogenetics and evolution in R language. Bioinformatics, 20(2), 289–290. https://doi.org/10.1093/bioinformatics/btg412

Pimm, S. L., Jenkins, C. N., Abell, R., Brooks, T. M., Gittleman, J. L., Joppa, L. N., Raven, P. H., Roberts, C. M., & Sexton, J. O. (2014). The biodiversity of species and their rates of extinction, distribution, and protection. Science, 344(6187), 1246752. https://doi.org/10.1126/science.1246752

Powers, R. P., & Jetz, W. (2019). Global habitat loss and extinction risk of terrestrial vertebrates under future land-use-change scenarios. Nature Climate Change, 9(4), 323–329. https://doi.org/10.1038/s41558-019-0406-z

Pressey, R. L., Cabeza, M., Watts, M. E., Cowling, R. M., & Wilson, K. A. (2007). Conservation planning in a changing world. Trends in Ecology and Evolution, 22(11), 583–592. https://doi.org/10.1016/j.tree.2007.10.001

Smith, S. A., & Brown, J. W. (2018). Constructing a broadly inclusive seed plant phylogeny. American Journal of Botany, 105(3), 302–314. https://doi.org/10.1002/ajb2.1019

Sugimoto, N., Fukasawa, K., Asahara, A., Kasada, M., Matsuba, M., & Miyashita, T. (2022). Positive and negative effects of land abandonment on butterfly communities revealed by a hierarchical sampling design across climatic regions. Proceedings of the Royal Society B: Biological Sciences, 289(1971). https://doi.org/10.1098/rspb.2021.2222

Swets, J. A. (1988). Measuring the accuracy of diagnostic systems. Science, 240(4857), 1285–1293. https://doi.org/10.1126/science.3287615

Tikhonov, G., Duan, L., Abrego, N., Newell, G., White, M., Dunson, D., & Ovaskainen, O. (2020). Computationally efficient joint species distribution modeling of big spatial data. Ecology, 101(2), e02929. https://doi.org/10.1002/ecy.2929

Vehtari, A., Gelman, A., & Gabry, J. (2017). Practical Bayesian model evaluation using leave-one-out cross-validation and WAIC. Statistics and Computing, 27(5), 1413–1432. https://doi.org/10.1007/s11222-016-9696-4

Walker, B. E., Leão, T. C. C., Bachman, S. P., Bolam, F. C., & Nic Lughadha, E. (2020). Caution needed when predicting species threat status for conservation prioritization on a global scale. Frontiers in Plant Science, 11, 520. https://doi.org/10.3389/fpls.2020.00520

Watanabe, E., Saito, M. U., Hayashi, N., & Matsuda, H. (2014). Biodiversity hotspot analysis based on the extinction risk of vascular plant species in the Red Data Book of Japan. Japanese Journal of Conservation Ecology, 19(1), 53–66. https://doi.org/10.18960/hozen.19.1_53

Watanabe, S. (2010). Asymptotic equivalence of Bayes cross validation and widely applicable information criterion in singular learning theory. Journal of Machine Learning Research, 11, 3571–3594.

Zhang, H. (2004). Inconsistent estimation and asymptotically equal interpolations in model-based geostatistics. Journal of the American Statistical Association, 99(465), 250–261. https://doi.org/10.1198/016214504000000241

Zhang, L., Datta, A., & Banerjee, S. (2019). Practical Bayesian modeling and inference for massive spatial datasets on modest computing environments. Statistical Analysis and Data Mining, 12(3), 197–209. https://doi.org/10.1002/sam.11413

